# Gmo labeling in food products in montevideo, uruguay

**DOI:** 10.1101/2020.08.17.254243

**Authors:** M. Arleo, P. Benavente, V. Cardozo, A. Katz, S. Vázquez, A. Da Silva Tavares, M. Legnani, C. Martínez Debat

**Affiliations:** Laboratorio de Trazabilidad Molecular Alimentaria, Sección Bioquímica, Facultad de Ciencias, Universidad de la República; Servicio de Regulación Alimentaria-Laboratorio de Bromatología, Intendencia de Montevideo; Núcleo Interdisciplinario Colectivo TÁ, Espacio Interdisciplinario, Universidad de la República

**Keywords:** GMOs, labeling, food legislation, consumer information, molecular techniques

## Abstract

Montevideo establishes the mandatory labeling of foods containing genetically modified material through the Departmental Decree No. 36.554, positioning Uruguay within the 65 countries that have incorporated this type of regulation.

The Food Regulation Service, in its role of sanitary police, and through its Laboratory of Bromatology, in agreement with the Food Molecular Traceability Laboratory (Faculty of Sciences, University of the Republic), carried out the analysis of 206 products made with ingredients derived from corn and/or soybean, during the 2015-2017 period, within the framework of compliance with the aforementioned Decree.

The strategy used consisted of the application of molecular techniques (Real Time PCR), for the detection of common sequences present in the transgenic events of soybean and corn, and the subsequent quantification of the content of GM material, in relation to an established labeling threshold of 1%.

As a result of this study, it was found that 36.9% of the analyzed foods presented sequences derived from genetically modified plant organisms (GMOs); and in 95% of the cases, its content exceeded the threshold established for its labeling.

This study, constitutes the first approach to the knowledge of different transgenic elements distribution in food commercialized in Montevideo.

These results provide valuable information to both the consumer, for decision making about the food to be consumed, and also for the official control organizations, which must enforce the regulations.

This type of work has demonstrated, once again, the importance of the interrelation between academy and state agencies, in the generation of knowledge and in the implementation of new analytical methodologies, as well as in the training of qualified human resources and in the compliance with current regulations.

## 1. Introduction

According to the World Health Organization (WHO), “genetic modified food” are those derived from organisms whose genetic material (DNA) has been modified in a way that does not occur naturally, for example, through the introduction of a gene from a different organism (WHO, 2015).

In order to preserve the right to choose the consumption of genetically modified foods,, more than 60 countries have enacted legal measures to regulate their labeling; however, the specific characteristics of these regulations differ considerably from one country to another (Abad R. et al, 2004; Hilbeck et al., 2015; Dizon et al., 2016). Certain countries require mandatory labeling of foods that contain, are, or are derived from Genetically Modified Organisms (GMOs) (European Commission, 2003a, European Commission 2003b) while others have chosen to promote voluntary labeling (Acosta et al., 2015; Bovay, et. al, 2016; Just Label It Campaign, 2017). Despite the differences in these provisions, there is a broad consensus that the general objective of labeling is to inform consumers (Kamle et al., 2013). Also, it is assumed that such labeling is not a substitute for risk assessment of genetically modified food safety, but it serves as an additional and complementary control, in the regulatory process (J. Žel et al., 2012).

Uruguay is the tenth country with the highest production of transgenic crops worldwide and the fourth in South America after Brazil, Argentina and Paraguay (ISAAA, 2017). Since its introduction in 2003, ten transgenic varieties of corn, and five of soy, with agronomic characteristics of insects resistance and/or herbicide tolerance have been authorized for cultivation and processing. Also, thirteen additional soy and/or maize stacked events, which show combined agronomic characteristics for insect (and/or abiotic stress) resistance and/or insecticide tolerance, are in evaluation stages (Table 1)(GNBio,2017).

**Table 1.** Transgenic varieties approved for cultivation, consumption and processing in Uruguay (2003-2017) (Taken from GNbio, 2017; Biosafety Clearing-House, 2017)

| Species / Event | Agronomic characteristics | Status in Uruguay | Molecular Marker |  |  |  |
| --- | --- | --- | --- | --- | --- | --- |
|  |  |  | 35S | t-NOS | bar | FMV |
| *Glycine max* GTS 40-3-2 | Glyphosate tolerance | Approved 2/10/1996 | YES | YES | NO | NO |
| *G. max* A2704-12 (LL) | Ammonium glufosinate tolerance | Approved 19/9/2012 | YES | NO | NO | NO |
| *G. max* A5547-127 (LL) | Ammonium glufosinate tolerance | Approved 19/9/2012 | YES | NO | NO | NO |
| *G. max* M0N89788XM0N87701 (RR2YBt) | Glyphosate tolerance / Resistance to lepidóptera | Approved 19/9/2012 | NO | NO | NO | YES |
| *G. max* BPS-CV127-9 | Ammonium glufosinate tolerance | Approved 29/10/2014 | NO | NO | NO | NO |
| *G. max* DAS44406-6 | 2,4D tolerance | In evaluation | NO | NO | NO | NO |
| *G. max* MON89788XM0N87708 | Glyphosate tolerance | In evaluation | NO | NO | NO | YES |
| *G. max* FG72 | Glyphosate and HPPD inhibitor herbicides tolerance | In evaluation | NO | YES | NO | NO |
| *G. max* FG72XA5547-127 | Glyphosate, ammonium glufosinate and HPPD inhibitor herbicides tolerance | In evaluation | YES | YES | NO | NO |
| *G. max* HB4-(PAT) | Abiotic stress (drought) tolerance | In evaluation | NO | YES | NO | NO |
| *G. max* DAS44406-6XDAS81419-2 | 2,4D, glyphosate and ammonium glufosinate tolerance / Resistance to lepidoptera | In evaluation | NO | NO | NO | NO |
| *G. max* MON89788XM0N87701XM0N87708XM0N87751 | Dicamba and glyphosate tolerance / Resistance to lepidóptera | In evaluation | NO | NO | NO | YES |
| *Zea mays* MON810 | Resistance to lepidóptera | Approved 20/06/2003, 21/06/2011 | YES | NO | NO | NO |
| *Z. mays* BT11 | Resistance to lepidoptera / Ammonium glufosinate tolerance | Approved 05/05/2004, 21/06/2011 | YES | YES | NO | NO |
| *Z. mays* GA21 | Glyphosate tolerance | Approved 21/6/2011 | NO | YES | NO | NO |
| *Z. mays* GA21XBT11 | Glyphosate and ammonium glufosinate tolerance /Resistance to lepidoptera | Approved 21/6/2011 | YES | YES | NO | NO |
| *Z. mays* TC1507 | Resistance to lepidoptera / Ammonium glufosinate tolerance | Approved 21/06/11, 20/10/11 | YES | NO | NO | NO |
| *Z. mays* NK603 | Glyphosate tolerance | Approved 21/6/2011 | YES | YES | NO | NO |
| *Z. mays* M0N810XNK603 | Glyphosate tolerance / Resistance to lepidóptera | Approved 21/6/2011 | YES | YES | NO | NO |
| *Z. mays* TC1507XNK603 | Resistance to lepidoptera / Glyphosate and ammonium glufosinate tolerance | Approved 19/9/2012 | YES | YES | NO | NO |
| *Z. mays* BT11XMIR162XGA21 (*) | Resistance to lepidoptera / Glyphosate and ammonium glufosinate tolerance | Approved 21/9/2012 | YES | YES | NO | NO |
| *Z. mays* M0N89034XTC1507XNK603 (*) | Glyphosate and ammonium glufosinate tolerance / Resistance to lepidoptera | Approved 21/9/2012 | YES | YES | NO | NO |
| *Z. mays* M0N89034XM0N88017 | Glyphosate tolerance /Resistance to lepidoptera and coleóptera | In evaluation | YES | YES | NO | NO |
| *Z. mays* BT11XMIR162XMIR604XGA21 | Glyphosate and ammonium glufosinate tolerance / Resistance to lepidoptera and coleóptera | In evaluation | YES | YES | NO | NO |
| *Z. mays* MON89034XNK603XTC1507XDAS40278-9 | 2,4D, glyphosate and ammonium glufosinate tolerance / Resistance to lepidoptera | In evaluation | YES | YES | NO | NO |
| *Z. mays* MON810XTC1507XNK603 | Resistance to lepidoptera / Glyphosate and ammonium glufosinate tolerance | In evaluation | YES | YES | NO | NO |
| *Z. mays* T25 | Ammonium glufosinate tolerance | In evaluation | YES | NO | NO | NO |

Uruguay regulates the use of genetically modified (GM) plants and their parts through a National Decree enacted in 2008. It promotes actions aimed at the implementation of the voluntary labeling “GM” or “non-GM”, applicable to those foods in which the presence of DNA or genetically modified proteins can be verified through final product analysis (Decree 353/2008, 2008). More recently, in 2015, Montevideo (the country’s capital where more than 50% of the Uruguayan population lives) decreed mandatory labeling of all foods containing genetically modified organisms in a percentage higher than a threshold of 1%. (Decrees N° 34.901, 2013; N° 35.099, 2014; and N°36.554, 2018). While this requirement applies only in Montevideo, it has prompted other three (out of 19) Uruguayan Departments to approve similar Decrees, and also a Decree of national scope is under discussion in the Uruguayan Parliament (Anzalone P., 2016) Given that in Uruguay most of soybean and corn crops are transgenic (ISAAA, 2017) it is possible that food made from raw materials derived from these species, contain genetically modified material, and that their presence is found in percentages that exceed the 1% threshold set in the regulation. In this context of GMOs regulation in Uruguay, it is essential to have analytical tools that guarantee compliance with the current Decrees.

Numerous methods have been described to detect GMO-derived material in food, feed and seeds (Fernandez, S., et al, 2005; Žel, J. et al., 2008; Querci, M. et al., 2010; Van den Eede et al., 2011; European Network of GMO laboratories (ENGL) 2011; Bonfini et al, 2012; Fraiture, M. A. et al., 2015), however, the most used methodology for this purpose is based on the detection of transgenic DNA using the Real Time PCR technique. (Van den Bulcke, M. et al., 2010; Barbau-Piednoir E et al., 2010; Cottenet, G. et al., 2013; Wu, Y., et al 2014; Huber, I. et al., 2013; Barbau-Piednoir E. et al., 2014; Bhoge, R. K. et al., 2016). Most studies in the available literature focuses on the design and development of these methods, but only a few study the GMO content in food found in the market (Cardarelli et.al, 2005; Viljoen, C. D., 2006; Prins, T. W.,et al, 2016; Elsanhoty et al., 2013).

The strategy for detecting and quantifying GMOs in food consists of several steps, as described in international standards ISO 24276: 2006, ISO 21569: 2005, ISO 21570: 2005, ISO 21571. First, DNA present in food is extracted; secondly, common DNA sequences sherd by most genetically modified plants are screened; in the case of a positive result, quantification of GMO content is performed.

The present study describes the use of molecular methods, based on Real Time PCR, for the detection, identification and quantification of genetically modified DNA sequences in foods manufactured from corn and soybean derived ingredients, sampled in Montevideo Department, between 2015 and 2017. This study was carried out within the framework of a collaboration and technological transfer Legal Agreement between the Official Service of Food Regulation of the Montevideo Municipality (Laboratory of Bromatology) and the Academy (Food Molecular Traceability Laboratory, Faculty of Sciences, University of the Republic).

## 2. Materials and Methods

### 2.1 Sampling

Between 2015 and 2017, the regulatory agency (Laboratory of Bromatology of the Municipality of Montevideo) conducted the sampling of different foods made with ingredients derived from corn or soybean (according to information declared by the manufacturer), from different commercial stores in the capital. These were chosen taking into account that in Uruguay only some transgenic lines of maize and soya species are authorized for consumption and processing (GNBio, 2017), so that only these two probably GM species should be present in commercialized foods.

Products containig both species were not included in this study, due to technical reasons. Products from different provenances, national and imported, were sampled, including those marketed by small producers and by multinational companies, provided that they did not present the label required by Decree 36.554.

Product sampling was done according to the strategy established in the international standard ISO 21568 for the detection of GMOs in foodstuff (ISO 21568, 2005).

### 2.2 DNA Purification

Genomic DNA was extracted from 200 milligrams of homogenized material obtained from the processing of five units of each sample. Commercial kits, validated for use in complex food matrices, and following the manufacturer’s specifications, were used. Foods with soy ingredients were processed using the *SureFood® PREP Advanced* kit (R-BIOPHARM), while foods made with corn were processed using the *foodproof GMO extraction* kit (BIOTECON Diagnostics). The obtained DNA was quantified by spectrophotometry using the NanoDrop 1000 micro-volume spectrophotometer (Thermo Scientific, USA).

### 2.3 Detection of transgenic DNA sequences

Real Time PCR technique was used to detect both the presence of plant DNA and four transgenicity DNA markers: the *35S* promoter sequence from Cauliflower Mosaic Virus (CaMV); the *Agrobacterium tumefasciens* terminator *nos* from nopaline-synthase enzyme (*t-NOS*); the *Scrophularia* mosaic virus promoter (*FMV*); and the bar gene that codes phosphinothricin acetyltransferase enzyme (*pat*), from *Streptomyces hygroscopicus*. These markers are found in a wide variety of genetically modified corn and soybean lines released worldwide (ISAAA, 2017, Biosafety Clearing-House 2017), as well as in those authorized for consumption and processing in Uruguay. (GNbio, 2017) (Table 1).

DNA amplification was carried out using commercial kits: *foodproof® GMO Screening Kit* (*35S, t-NOS, bar, FMV*) (BIOTECON Diagnostics) and *SureFood® GMO SCREEN 4plex 35S/NOS/FMV/IAC* (R-Biopharm), following manufacturers’ specifications. All reactions were performed using the ABI 7500 PCR System thermocycler from Applied Biosystems.

### 2.4 Quantification of corn and soybean GM DNA

For the quantification of DNA from GM maize, the commercial kit *foodproof® GMO 35S Maize Quantification Kit* (BIOTECON Diagnostics) was used, while for the quantification of GM soybean, the commercial kit *SureFood GMO Quant RR Soya* (R-BIOPHARM) was used. All amplifications were performed using the ABI 7500 PCR System thermocycler from Applied Biosystems.

### 2.5 Statistical analysis

To compare the results of the tests, Fisher’s exact test was used, under the null hypothesis that there are no significant differences between the variables (Fisher, R.A, 1954). In all cases, a probability level of p=0.05 was used to accept or reject the hypothesis. All statistical analysis were performed using online GraphPad program (http://www.graphpad.com/quickcalcs/contingency1/)

## 3 Results and discussion

### 3.1 Sampling of food coming from corn or soybean

A total of 206 food products (n = 206) made with ingredients derived from either corn (n = 128) or soybean (n = 78), were sampled for this study.

Sampled products were classified into two categories according to the degree of food processing: natural or minimally processed foods (processed, n = 72), and ultra-processed products (ultra-processed, n = 134), the latter group being understood as consisting of those foods that have gone through multiple processes of industrialization, and containing high amounts of additives, preservatives, stabilizers, and flavorings (Ministry of Public Health, 2016). (Table 2)

**Tabla 2.** Food analyzed in the 2015-2017 period. They were categorized based on product type (processed and ultra-processed foods), and according to the content of the vegetable species: corn or soybean, as stated in the market labeling.

| Food categories | Made with corn | Made with soybean | Total |
| --- | --- | --- | --- |
| **Processed** | **n=62** | **n=10** | **n= 72** |
| Natural or canned grains | 39 | 6 | 45 |
| Flours | 17 | - | 17 |
| Starches | 6 | - | 6 |
| Vegetal proteins | - | 4 | 4 |
| **Ultra-processed** | **n=66** | **n=68** | **n= 134** |
| Cookies | 12 | 29 | 41 |
| CopSnacks | 17 | 10 | 27 |
| Deli Meats | - | 23 | 23 |
| Cereals | 22 | - | 22 |
| Sauces and juices | 7 | 6 | 13 |
| Dry pasta | 8 | - | 8 |

### 3.2 GM sequences detection

The results of this study reveal that various foods made with ingredients derived from soybeans and corn, have DNA sequences from plant genetically modified organisms. These sequences were found in 36.9% of the foods analyzed in this study (76/206), being found in 29.7% of foods prepared with corn (38/128), and in 48.7% of products made with soybean (38/78), with a statistically significant difference between the two matrices (p=0,0074) (Table 3).

**Table 3.**
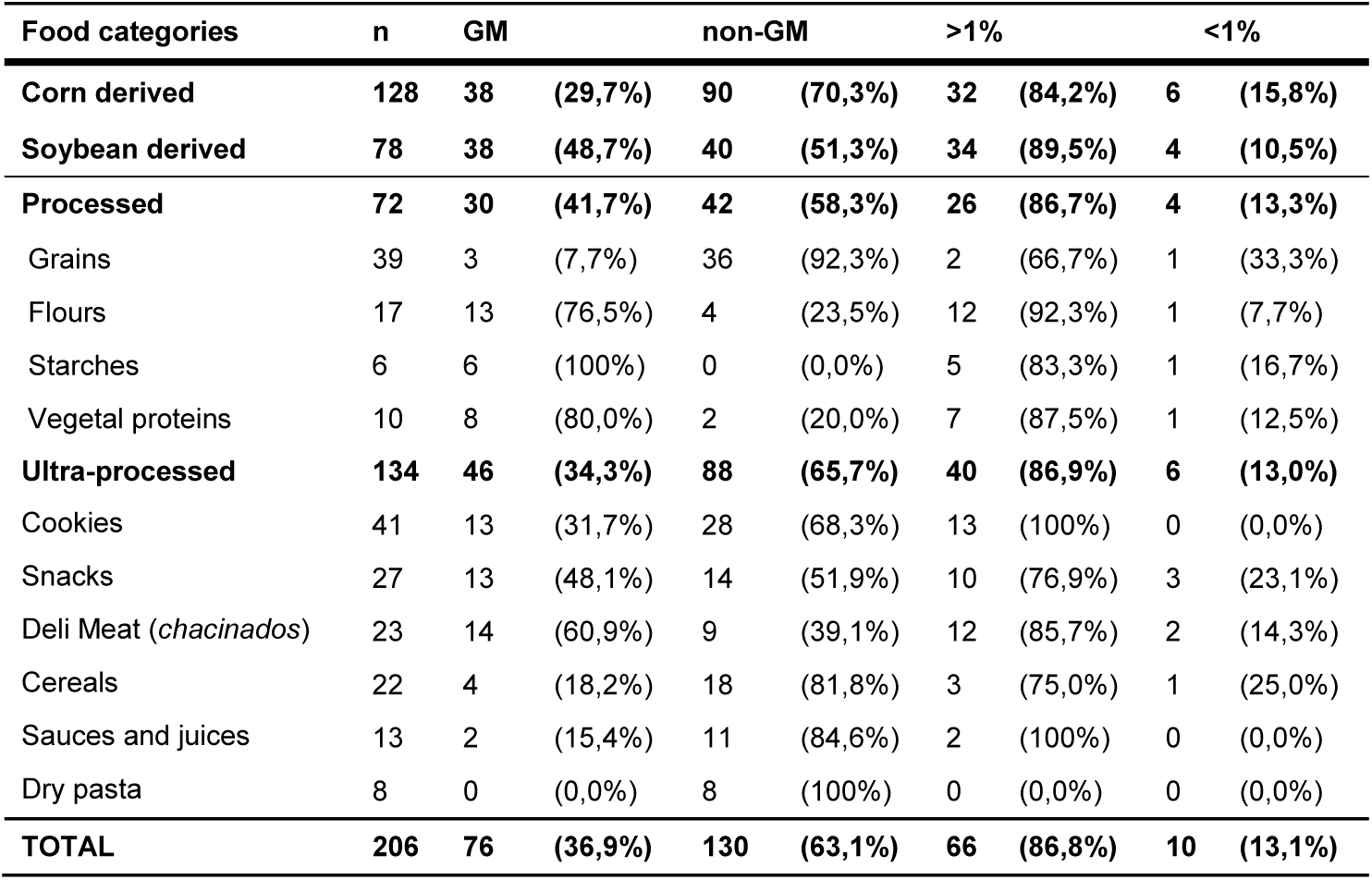
Results of the analysis of transgenic sequences detection in foods made with ingredients derived from corn or soybeans. “GM”: positive samples for the presence of transgenic sequences. “Non-GM”: negative samples for the presence of transgenic sequences. “> 1%”: content of GM material greater than 1%. “<1%”: content of GM material less than 1%.

On the other hand, the frequencies of presence of GM sequences in the processed and ultra-processed products were similar, without significant differences (p=0.3637) (Table 3), unlike that reported in other related study. Gonzales and collaborators, found a lower GM maize presence in Mexican Artisan products, in comparison to Industrial Tortillas. This difference between the studies could be due to a difference in the sample universe, and/or also in the type and number of samples taken in both cases (Gonzales et al, 2017).

Among processed products derived from corn, starch showed a presence of 100% transgenes (6/6), vegetable protein 80% (8/10), and corn flour 76.5% (13/17). The natural grains showed a presence of GM sequences of 7.7% (3/39). Among the ultra processed products, the deli meat products (*chacinados*) showed a frequency of transgene presence 60.9% (14/23), whereas snacks 48.1% (13/27). Dry pasta showed no presence of GM sequences (0/8).

It was also observed that the 35S promoter sequence was present in all the samples that were positive for the detection test of GM material (76/76), while the t-NOS marker was found in 94.7% of them (72/76) (Table 4). Furthermore, the FMV promoter sequence was found in 51.3% of GM positive samples (39/76), showing a higher frequency in the products made with soybean when compared to those made with corn (65.8% versus 36.8% in corn products). The sequence of the bar gene was not present in any of the analyzed products (Table 4).

**Table 4.** Transgenic sequences in processed and ultra-processed foods made with ingredients derived from corn or soybeans.

| Food categories | GM | p35S | t-NOS | FMV | bar |
| --- | --- | --- | --- | --- | --- |
| **Corn** | **38** | **38 (100%)** | **37 (97.4%)** | **14 (36.8%)** | **0 (0.0%)** |
| **Soybean** | **38** | **38 (100%)** | **35 (92.1%)** | **25 (65.8%)** | **NA NA** |
| **Processed** | **30** | **30 (100%)** | **29 (96.6%)** | **11 (36.7%)** | **0 (0.0%)** |
| **Ultra- processed** | **46** | **46 (100%)** | **43 (93.5%)** | **28 (60.9%)** | **0 (0.0%)** |
| **TOTAL** | **76** | **76 (100%)** | **72 (94.7%)** | **39 (51.3%)** | **0 (0.0%)** |

Of all products tested, only two (2/76) showed a single transgenic element, this being the p35S. In no case was the unique presence of *t-NOS* or *FMV* sequences observed. At least two or three of these elements were present in 94.7% (72/76) of the samples. The sequences *p35S* and *t-NOS* were both present in 46.0% (35/76) of the GM positive samples,, and the combination *p35S, t-NOS* and *FMV* was found in 48,7% (37/76) of them (not shown).

The high frequency of appearance of *35S* promoter and *NOS* terminator sequences in the analyzed samples, is in agreement with those described in the available bibliography, where it is described that both sequences are present in more than 80% of the transgenic events released and authorized for consumption, worldwide (ISAAA, 2017; Biosafety Clearing-House 2017). It should be noted that in Uruguay (and also in Southamerica), the transgenic varieties that are grown to a greater extent, (Bt11 and NK603 corn events, and GTS-40-3-2 soybean event) contain these two sequences in their genic construction (INASE, 2017). It was also observed that two of the analyzed products (a ground maize and a certain variety of sweet cookies) presented the *35S* promoter as the only transgenic sequence, which could be explained by the presence of some GM variety that does not contain the NOS terminator, such as the Mon810 corn event, also grown in our country and in the region (GNBio, 2017). Furthermore, the *FMV* promoter sequence was found in 51.3% of cases, always with *p35S* and *t-NOS* sequences. This genetic element is usually found in a few soybean events, for example in the MON89788 variety (RR2Y, authorized in Uruguay for production of export seeds), in the variety MON89788XMIN87701 (RR2YBt, authorized for consumption), and also in the MON87705 variety, not yet authorized in our country. Also, this element is found in a single variety of corn, MON89034, which in Uruguay is authorized for consumption in the form of the MON89034XTC1507XNK603 stacked event. It should be noted that the sequence of the *bar* gene was not present in any of the analyzed products. This result is consistent with the fact that to date, in the world, only 7 corn and 2 soybean events have been authorized that contain this transgene, and they are grown in the United States, Japan and the Philippines. Only one of these events, the Bt176 maize variety iscommerciallyauthorized in Argentina (Biosafety Clearing-House 2017). The low frequency of the *bar* gene presence, and that of Bt176 event in mass consumption foods made from corn has already been reported in similar studies (Dinon et al, 2010; Gonzales Ortega et al, 2017).

### 3.4 GMO Quantification

The quantification analysis of genetically modified material revealed that 86.8% (66/76) of the foods that were positive for detection of transgene sequences presented percentages greater than 1%, 84.2% of foods made from corn, and 89.5% of foods made from soy were found in this situation. Only 13.1% of the products that presented transgenic sequences (10/76), showed a content of GM material below this threshold. Among these, 4 were processed products and 6 were ultra-processed (Table 3).

The presence of GM material below the labeling threshold could be explained either by the use of different non-GM and GM ingredients that result in a final blend with low levels of GM material, or by unintentional contamination in the production chain. The latter could be the consequence oftransgene flow in crop production, contamination of the seeds during transportation, or during ingredients processing in the manufacturing plant (Aung et al, 2014).

It is important to note that although the legislation on the labeling of foods containing or derived from GMOs is in force in Montevideo since 2013, only a small fraction of foods made from corn and soybeans were found labeled, and these were not included in this study.

On the other hand, several products, national and imported, were observed with the legend “Non-transgenic certificate”, “GMO free” or “Non GMO”. This type of labeling is regulated in some countries such as the United States, where voluntary labeling is allowed for the presence and absence of GMOs (Albert, J. 2010; Just Label It, 2017). However, in Uruguay, unlike its capital Montevideo, there is still no regulation that contemplates or requires this distinction in the products. Among the foods labeled as “Non GMO”, there is a soy sauce, two cereal mixes, a deli meats and a vegetable protein. Analysis of these products revealed the presence of transgenes in two of these samples.

## 4. Conclusions

This study evidenced the presence of sequences derived from genetically modified plant organisms in 36.9% of the foods analyzed, marketed in Montevideo. In addition, 95% of these foods showed a content exceeding the threshold percentage set for labeling GM material

These results allow to affirm that there is a greater presence of GM material in products that contain soybean compared to those that contain corn. Likewise, it was found that the contents of GM material between the processed and ultra-processed products were similar.

To date this study, constitutes the first approach to the knowledge of the distribution of different transgenic elements in certain products that are commercialized in Montevideo. This type of study highlights the need for policies regarding the labeling of foods derived from GMOs, in order to guarantee the consumer’s right to know. Similarly, it provides objective information on the composition of foods, and enhances the importance of creating an effective regulatory system for monitoring and controlling this type of foods.

It would be interesting in the future to search for not yet authorized corn and soybean events in Uruguay, as well as analyzing new transgenic varieties of other plant species likely to be present in food. Although several uruguayan Departments have already chosen to incorporate the labeling of GM material in food, it would be possible to consider the formulation of a law with national scope.

## Supplementary Material

**Table I.** Food made from soybeans or corn, domestic and imported. Specific brands are omitted, and local typical names are shown in italics. Results of detection of: *p35:* CaMV *35S* promoter; *t-NOS:* Nos terminator; *FMV*: *Scrophularia* mosaic virus promoter; *bar*: gene that codes phosphinothricin acetyltransferase enzyme from *Streptomyces hygroscopicus*. NA: Not assayed. NL: No GMO labeling

| Muestra | Product type (in label) | Category | Species | Country of origin | GM Screening | *p35s* | *t-NOS* | *FMV* | *bar* | Quantification | Labeling |
| --- | --- | --- | --- | --- | --- | --- | --- | --- | --- | --- | --- |
| 1 | Polenta corn flour | Processed | Corn | ARGENTINA | + | + | - | - | - | > 1% | NL |
| 2 | Polenta corn flour | Processed | Corn | URUGUAY | + | + | + | - | - | > 1% | NL |
| 3 | Polenta corn flour | Processed | Corn | URUGUAY | + | + | + | - | - | > 1% | NL |
| 4 | Polenta corn flour | Processed | Corn | URUGUAY | + | + | + | - | - | > 1% | NL |
| 5 | Polenta corn flour | Processed | Corn | URUGUAY | + | + | + | - | - | > 1% | NL |
| 6 | Polenta corn flour | Processed | Corn | URUGUAY | + | + | + | - | - | > 1% | NL |
| 7 | Polenta corn flour | Processed | Corn | URUGUAY | + | + | + | - | - | > 1% | NL |
| 8 | Polenta corn flour | Processed | Corn | URUGUAY | + | + | + | - | - | > 1% | NL |
| 9 | Polenta corn flour | Processed | Corn | URUGUAY | + | + | + | - | - | > 1% | NL |
| 10 | Polenta corn flour | Processed | Corn | URUGUAY | + | + | + | - | - | > 1% | NL |
| 11 | Instant biological polenta | Processed | Corn | ITALIA | - | - | - | - | - | NA | NL |
| 12 | Precooked corn flour for polenta | Processed | Corn | ARGENTINA | + | + | + | + | - | <1% | NL |
| 13 | Corn flour to prepare polenta | Processed | Corn | URUGUAY | - | - | - | - | - | NA | NL |
| 14 | Corn flour to prepare polenta | Processed | Corn | ITALIA | - | - | - | - | - | NA | NL |
| 15 | Precooked corn flour with cheese | Processed | Corn | ITALIA | - | - | - | - | - | NA | NL |
| 16 | Corn flour | Processed | Corn | URUGUAY | + | + | + | + | - | > 1% | NL |
| 17 | Corn flour to prepare polenta | Processed | Corn | URUGUAY | + | + | + | - | - | > 1% | NL |
| 18 | Cornstarch | Processed | Corn | ARGENTINA | + | + | + | + | - | > 1% | NL |
| 19 | Cornstarch | Processed | Corn | BRASIL | + | + | + | - | - | > 1% | NL |
| 20 | Cornstarch | Processed | Corn | URUGUAY | + | + | + | - | - | > 1% | NL |
| 21 | Gluten-free Cornstarch | Processed | Corn | ARGENTINA | + | + | + | + | - | > 1% | NL |
| 22 | Cornstarch | Processed | Corn | URUGUAY | + | + | + | + | - | > 1% | NL |
| 23 | Wheat semolina | Processed | Corn | URUGUAY | + | + | + | - | - | <1% | NL |
| 24 | Yellow shelled corn | Processed | Corn | BRASIL | - | - | - | - | - | NA | NL |
| 25 | Natural grain corn | Processed | Corn | BRASIL | - | - | - | - | - | NA | NL |
| 26 | Sweet corn grain without added salt | Processed | Corn | EEUU | - | - | - | - | - | NA | NL |
| 27 | Whole yellow corn kernels | Processed | Corn | ARGENTINA | - | - | - | - | - | NA | NL |
| 28 | Whole yellow corn kernels | Processed | Corn | TAILANDIA | - | - | - | - | - | NA | NL |
| 29 | Whole yellow corn kernels | Processed | Corn | FRANCIA | - | - | - | - | - | NA | NL |
| 30 | Corn kernels natural | Processed | Corn | BRASIL | - | - | - | - | - | NA | NL |
| 31 | Corn kernels natural | Processed | Corn | BRASIL | - | - | - | - | - | NA | NL |
| 32 | Whole grain corn | Processed | Corn | CHINA | - | - | - | - | - | NA | NL |
| 33 | Sweet corn in vacuum packed grains | Processed | Corn | FRANCIA | - | - | - | - | - | NA | NL |
| 34 | Creamy yellow corn | Processed | Corn | ARGENTINA | - | - | - | - | - | NA | NL |
| 35 | Steamed sweet corn | Processed | Corn | BRASIL | - | - | - | - | - | NA | NL |
| 36 | Shelled sweet corn | Processed | Corn | URUGUAY | - | - | - | - | - | NA | NL |
| 37 | Natural sweet corn | Processed | Corn | FRANCIA | - | - | - | - | - | NA | NL |
| 38 | Corn, carrots and peas | Processed | Corn | CHINA | - | - | - | - | - | NA | NL |
| 39 | Mixed vegetables | Processed | Corn | EEUU | - | - | - | - | - | NA | NL |
| 40 | Jardiniere of vegetables and legumes | Processed | Corn | ARGENTINA | - | - | - | - | - | NA | NL |
| 41 | Corn and peas | Processed | Corn | BRASIL | - | - | - | - | - | NA | NL |
| 42 | Canned peas and corn | Processed | Corn | BRASIL | - | - | - | - | - | NA | NL |
| 43 | Peas and corn | Processed | Corn | BRASIL | - | - | - | - | - | NA | NL |
| 44 | Corn cream | Processed | Corn | BRASIL | - | - | - | - | - | NA | NL |
| 45 | Corn cream | Processed | Corn | TAILANDIA | - | - | - | - | - | NA | NL |
| 46 | Corn kernels natural | Processed | Corn | BRASIL | - | - | - | - | - | NA | NL |
| 47 | Corn kernels natural | Processed | Corn | BRASIL | - | - | - | - | - | NA | NL |
| 48 | Corn kernels natural | Processed | Corn | BRASIL | - | - | - | - | - | NA | NL |
| 49 | Rehydrated peas and corn grain | Processed | Corn | BRASIL | - | - | - | - | - | NA | NL |
| 50 | Whole grain corn | Processed | Corn | BRASIL | - | - | - | - | - | NA | NL |
| 51 | Corn kernels natural | Processed | Corn | BRASIL | - | - | - | - | - | NA | NL |
| 52 | Jardiniere of vegetables and legumes | Processed | Corn | ARGENTINA | + | + | + | - | - | <1% | NL |
| 53 | *Pisingallo* corn | Processed | Corn | ARGENTINA | - | - | - | - | - | NA | NL |
| 54 | Trod corn to make porridge | Processed | Corn | URUGUAY | + | + | + | - | - | > 1% | NL |
| 55 | Corn pisingallo to make popcorn | Processed | Corn | ARGENTINA | - | - | - | - | - | NA | NL |
| 56 | Caramelized Popcorn | Processed | Corn | URUGUAY | - | - | - | - | - | NA | NL |
| 57 | Sweet popcorn | Processed | Corn | URUGUAY | - | - | - | - | - | NA | NL |
| 58 | *Pisingallo* corn | Processed | Corn | URUGUAY | - | - | - | - | - | NA | NL |
| 59 | *Pisingallo* corn | Processed | Corn | ARGENTINA | - | - | - | - | - | NA | NL |
| 60 | Scarlet *mazamorra* | Processed | Corn | ARGENTINA | + | + | + | + | - | > 1% | NL |
| 61 | Mixture to prepare popcorn | Processed | Corn | BRASIL | - | - | - | - | - | NA | NL |
| 62 | Popcorn Corn | Processed | Corn | BRASIL | - | - | - | - | - | NA | NL |
| 63 | Corn gluten-free pasta | Ultra-Processed | Corn | RUMANIA | - | - | - | - | - | NA | NL |
| 64 | Corn gluten-free pasta | Ultra-Processed | Corn | RUMANIA | - | - | - | - | - | NA | NL |
| 65 | Corn gluten-free pasta | Ultra-Processed | Corn | AUSTRALIA | - | - | - | - | - | NA | NL |
| 66 | Corn gluten-free pasta | Ultra-Processed | Corn | RUMANIA | - | - | - | - | - | NA | NL |
| 67 | White and yellow corn paste, and rice | Ultra-Processed | Corn | ITALIA | - | - | - | - | - | NA | NL |
| 68 | Dry pasta Italian spaghetti type | Ultra-Processed | Corn | AUSTRALIA | - | - | - | - | - | NA | NL |
| 69 | Dry pasta with garlic and parsley flavor | Ultra-Processed | Corn | ITALIA | - | - | - | - | - | NA | NL |
| 70 | Dry pasta with vegetables | Ultra-Processed | Corn | AUSTRALIA | - | - | - | - | - | NA | NL |
| 71 | Nutricereal with honey and oats | Processed | Corn | URUGUAY | + | + | + | - | - | > 1% | NL |
| 72 | Granola assortment of seeds with honey | Processed | Corn | URUGUAY | - | - | - | - | - | NA | NL |
| 73 | Cereals | Ultra-Processed | Corn | URUGUAY | + | + | + | - | - | > 1% | NL |
| 74 | Cornflakes | Ultra-Processed | Corn | ARGENTINA | - | - | - | - | - | NA | NL |
| 75 | Cornflakes | Ultra-Processed | Corn | ARGENTINA | - | - | - | - | - | NA | NL |
| 76 | Cornflakes | Ultra-Processed | Corn | ARGENTINA | - | - | - | - | - | NA | NL |
| 77 | Alimento estrusado | Ultra-Processed | Corn | ARGENTINA | - | - | - | - | - | NA | NL |
| 78 | Alimento a base de cereales | Ultra-Processed | Corn | BRASIL | + | + | + | + | - | > 1% | NL |
| 79 | Sweetened cornflakes | Ultra-Processed | Corn | ARGENTINA | - | - | - | - | - | NA | NL |
| 80 | Sweetened cornflakes | Ultra-Processed | Corn | EEUU | - | - | - | - | - | NA | "Non GMO" |
| 81 | Cornflakes | Ultra-Processed | Corn | EEUU | - | - | - | - | - | NA | "Non GMO" |
| 82 | Cornflakes | Ultra-Processed | Corn | ARGENTINA | - | - | - | - | - | NA | NL |
| 83 | Natural cornflakes | Ultra-Processed | Corn | ARGENTINA | - | - | - | - | - | NA | NL |
| 84 | Dietetic cereals | Ultra-Processed | Corn | CHILE | - | - | - | - | - | NA | NL |
| 85 | Granola with nuts | Processed | Corn | URUGUAY | - | - | - | - | - | NA | NL |
| 86 | Food based on corn flour | Processed | Corn | ARGENTINA | + | + | + | + | - | <1% | NL |
| 87 | Sweetened cornflakes | Ultra-Processed | Corn | BRASIL | - | - | - | - | - | NA | NL |
| 88 | Dietetic food with added vitamins | Ultra-Processed | Corn | BRASIL | - | - | - | - | - | NA | NL |
| 89 | Cereals and dehydrated fruits mix | Processed | Corn | ALEMANIA | - | - | - | - | - | NA | NL |
| 90 | Cornflakes | Ultra-Processed | Corn | BRASIL | - | - | - | - | - | NA | NL |
| 91 | Cornflakes | Ultra-Processed | Corn | FRANCIA | - | - | - | - | - | NA | NL |
| 92 | Food based on cereals, honey, nuts | Processed | Corn | URUGUAY | - | - | - | - | - | NA | "No transgénico" |
| 93 | Tortillas wraps | Ultra-Processed | Corn | ESPAÑA | - | - | - | - | - | NA | NL |
| 94 | Corn-based snack | Ultra-Processed | Corn | MEXICO | - | - | - | - | - | NA | NL |
| 95 | Corn-based snack with cheese flavor | Ultra-Processed | Corn | EEUU | + | + | + | + | - | > 1% | NL |
| 96 | Corn-based snack with cheese flavor | Ultra-Processed | Corn | BÉLGICA | - | - | - | - | - | NA | NL |
| 97 | Tortillas | Ultra-Processed | Corn | ALEMANIA | - | - | - | - | - | NA | NL |
| 98 | Baked corn toast | Ultra-Processed | Corn | MEXICO | + | + | + | - | - | <1% | NL |
| 99 | Salted Corn Fried Snack Product | Ultra-Processed | Corn | BÉLGICA | - | - | - | - | - | NA | NL |
| 100 | Gluten free corn toast | Ultra-Processed | Corn | AUSTRALIA | - | - | - | - | - | NA | NL |
| 101 | Sticks with ham flavor | Ultra-Processed | Corn | ARGENTINA | - | - | - | - | - | NA | NL |
| 102 | Corn flour sticks with cheese flavor | Ultra-Processed | Corn | EEUU | + | + | + | + | - | > 1% | NL |
| 103 | Corn Product corn based | Ultra-Processed | Corn | ARGENTINA | + | + | + | + | - | > 1% | NL |
| 104 | Tacos dough | Ultra-Processed | Corn | EEUU | + | + | + | - | - | <1% | NL |
| 105 | Roast beef flavor snack | Ultra-Processed | Corn | ARGENTINA | + | + | + | - | - | > 1% | NL |
| 106 | Vegatable mix flavor snack | Ultra-Processed | Corn | ITALIA | - | - | - | - | - | NA | NL |
| 107 | Cheese flavor corn butty | Ultra-Processed | Corn | URUGUAY | + | + | + | - | - | <1% | NL |
| 108 | Quinoa based snack product | Ultra-Processed | Corn | REINO UNIDO | - | - | - | - | - | NA | NL |
| 109 | Corn *tortilla* | Ultra-Processed | Corn | URUGUAY | + | + | + | + | - | > 1% | NL |
| 110 | Cornmeal and cocoa cookies with chocolate | Ultra-Processed | Corn | ESPAÑA | - | - | - | - | - | NA | NL |
| 111 | Powder to prepare vanilla flavor muffins | Ultra-Processed | Corn | URUGUAY | + | + | + | + | - | > 1% | NL |
| 112 | Powder to prepare sponge cake | Ultra-Processed | Corn | URUGUAY | - | - | - | - | - | NA | NL |
| 113 | Sweet cookies with bran | Ultra-Processed | Corn | ARGENTINA | - | - | - | - | - | NA | NL |
| 114 | Lemon flavor pudding | Ultra-Processed | Corn | ARGENTINA | - | - | - | - | - | NA | NL |
| 115 | Chocolate and vanilla flavor pudding | Ultra-Processed | Corn | URUGUAY | - | - | - | - | - | NA | NL |
| 116 | Vanilla flavor dessert powder | Ultra-Processed | Corn | URUGUAY | - | - | - | - | - | NA | NL |
| 117 | Vanilla flavor dessert powder | Ultra-Processed | Corn | URUGUAY | - | - | - | - | - | NA | NL |
| 118 | Sweet cookie *María* type without gluten | Ultra-Processed | Corn | ITALIA | - | - | - | - | - | NA | NL |
| 119 | Vanilla flavor dessert powder | Ultra-Processed | Corn | ESPAÑA | - | - | - | - | - | NA | NL |
| 120 | Corn cookies | Ultra-Processed | Corn | URUGUAY | + | + | + | - | - | > 1% | NL |
| 121 | Corn cookies | Ultra-Processed | Corn | URUGUAY | + | + | + | - | - | > 1% | NL |
| 122 | White sauce | Ultra-Processed | Corn | URUGUAY | + | + | + | + | - | > 1% | NL |
| 123 | Mayonnaise light | Ultra-Processed | Corn | ARGENTINA | - | - | - | - | - | NA | NL |
| 124 | Tartar sauce | Ultra-Processed | Corn | REINO UNIDO | - | - | - | - | - | NA | NL |
| 125 | Corn and butter instant cream soup | Ultra-Processed | Corn | ARGENTINA | - | - | - | - | - | NA | NL |
| 126 | Ketchup sauce reduced in caloric value | Ultra-Processed | Corn | BRASIL | - | - | - | - | - | NA | NL |
| 127 | Ketchup sauce | Ultra-Processed | Corn | URUGUAY | - | - | - | - | - | NA | NL |
| 128 | Mustard-based seasoning | Ultra-Processed | Corn | URUGUAY | - | - | - | - | - | NA | NL |
| 129 | Cooked peeled chorizo, *choripan* | Ultra-Processed | Soybean | URUGUAY | + | + | + | + | - | > 1% | "Soybean<br>certificada no<br>transgénica" |
| 130 | Milanese of chard, spinach and soy | Ultra-Processed | Soybean | ARGENTINA | + | + | + | + | - | > 1% | NL |
| 131 | Hotdogs light | Ultra-Processed | Soybean | URUGUAY | + | + | + | + | - | > 1% | NL |
| 132 | *Chorizo* flavored burger | Ultra-Processed | Soybean | URUGUAY | + | + | + | - | - | > 1% | NL |
| 133 | Beef burger | Ultra-Processed | Soybean | URUGUAY | - | - | - | - | - | NA | NL |
| 134 | Extra *chorizo* | Ultra-Processed | Soybean | URUGUAY | + | + | + | - | - | > 1% | NL |
| 135 | Chicken nuggets with cheese | Ultra-Processed | Soybean | BRASIL | - | - | - | - | - | NA | NL |
| 136 | Frankfurter type sausage | Ultra-Processed | Soybean | URUGUAY | + | + | + | + | - | > 1% | NL |
| 137 | Frankfurter type sausage | Ultra-Processed | Soybean | URUGUAY | + | + | + | + | - | > 1% | NL |
| 138 | *Longaniza* | Ultra-Processed | Soybean | URUGUAY | + | + | + | + | - | > 1% | NL |
| 139 | Beef burger | Ultra-Processed | Soybean | URUGUAY | + | + | + | + | - | > 1% | NL |
| 140 | Pasteurized *chorizo* | Ultra-Processed | Soybean | URUGUAY | - | - | - | - | - | NA | NL |
| 141 | Frankfurter type sausage | Ultra-Processed | Soybean | URUGUAY | + | + | + | + | - | > 1% | NL |
| 142 | Frankfurter type sausage | Ultra-Processed | Soybean | URUGUAY | + | + | + | + | - | > 1% | NL |
| 143 | Frozen meat burger | Ultra-Processed | Soybean | URUGUAY | + | + | + | + | - | <1% | NL |
| 144 | Frankfurter type sausage | Ultra-Processed | Soybean | URUGUAY | - | - | - | - | - | NA | NL |
| 145 | Extra *chorizo* with cheese and olives | Ultra-Processed | Soybean | URUGUAY | + | + | + | + | - | > 1% | NL |
| 146 | Low sodium super frozen burgers | Ultra-Processed | Soybean | URUGUAY | - | - | - | - | - | NA | NL |
| 147 | Frozen burgers | Ultra-Processed | Soybean | URUGUAY | + | + | + | + | - | <1% | NL |
| 148 | Seasoned and breaded ground chicken meat | Ultra-Processed | Soybean | BRASIL | - | - | - | - | - | NA | NL |
| 149 | Liver pate | Ultra-Processed | Soybean | BRASIL | - | - | - | - | - | NA | NL |
| 150 | Meat pate | Ultra-Processed | Soybean | BRASIL | - | - | - | - | - | NA | NL |
| 151 | Pork paste | Ultra-Processed | Soybean | FRANCIA | - | - | - | - | - | NA | NL |
| 152 | Soy milanese | Ultra-Processed | Soybean | ARGENTINA | + | + | + | + | - | > 1% | NL |
| 153 | Filet of soybean milanese type | Ultra-Processed | Soybean | URUGUAY | - | - | - | - | - | NA | NL |
| 154 | Mixture to prepare soybean steaks | Ultra-Processed | Soybean | URUGUAY | + | + | + | - | - | <1% | NL |
| 155 | Filet of soybean milanese type | Ultra-Processed | Soybean | URUGUAY | + | + | + | + | - | NA | "No transgénico" |
| 156 | Soybean protein textured and granulated | Processed | Soybean | ARGENTINA | + | + | + | + | - | > 1% | NL |
| 157 | Soybean protein granulated | Processed | Soybean | ARGENTINA | + | + | + | + | - | > 1% | NL |
| 158 | Soy beans | Processed | Soybean | URUGUAY | - | - | - | - | - | NA | NL |
| 159 | Soy beans | Processed | Soybean | URUGUAY | + | + | + | + | - | > 1% | NL |
| 160 | Soybean protein | Processed | Soybean | URUGUAY | + | + | + | - | - | > 1% | NL |
| 161 | Assortment of seeds with soy protein | Processed | Soybean | URUGUAY | + | + | + | - | - | > 1% | NL |
| 162 | Soy sauce | Ultra-Processed | Soybean | EEUU | - | - | - | - | - | NA | NL |
| 163 | Soy sauce | Ultra-Processed | Soybean | CHINA | - | - | - | - | - | NA | NL |
| 164 | Original soy sauce | Ultra-Processed | Soybean | EEUU | - | - | - | - | - | NA | NL |
| 165 | Vanilla flavor dessert powder | Ultra-Processed | Soybean | ARGENTINA | - | - | - | - | - | NA | NL |
| 166 | Sweet bread filled w/ sweet milk (*dulce de leche*) | Ultra-Processed | Soybean | URUGUAY | - | - | - | - | - | NA | NL |
| 167 | Vanilla flavor budin and chocolate chips | Ultra-Processed | Soybean | ARGENTINA | - | - | - | - | - | NA | NL |
| 168 | Premix for preparing rice bread | Ultra-Processed | Soybean | URUGUAY | + | + | + | + | - | > 1% | NL |
| 169 | Sweet piles stuffed with quince jam | Ultra-Processed | Soybean | ARGENTINA | + | + | + | + | - | > 1% | NL |
| 170 | Sweet cookies with oats, apples, and cinnamon | Ultra-Processed | Soybean | ARGENTINA | + | + | + | + | - | > 1% | NL |
| 171 | Sweet cookies | Ultra-Processed | Soybean | URUGUAY | + | + | - | - | - | > 1% | NL |
| 172 | Waffles filled with hazelnut paste | Ultra-Processed | Soybean | GRECIA | + | + | + | - | - | > 1% | NL |
| 173 | Sweet cake | Ultra-Processed | Soybean | BRASIL | + | + | + | - | - | > 1% | NL |
| 174 | Artisan wafers | Ultra-Processed | Soybean | URUGUAY | - | - | - | - | - | NA | NL |
| 175 | Sweet cookies decorated with chips | Ultra-Processed | Soybean | URUGUAY | + | + | + | - | - | > 1% | NL |
| 176 | Vanilla sponge cake with chocolate filling | Ultra-Processed | Soybean | POLONIA | - | - | - | - | - | NA | NL |
| 177 | Sugary biscuits | Ultra-Processed | Soybean | ARGENTINA | - | - | - | - | - | NA | NL |
| 178 | Sweet cookies filled with lemon flavor | Ultra-Processed | Soybean | ARGENTINA | - | - | - | - | - | NA | NL |
| 179 | Cookies with Milk | Ultra-Processed | Soybean | URUGUAY | - | - | - | - | - | NA | NL |
| 180 | Wafers with vanilla flavor filling | Ultra-Processed | Soybean | BRASIL | - | - | - | - | - | NA | NL |
| 181 | Waffles with lemon flavor filling | Ultra-Processed | Soybean | GRECIA | - | - | - | - | - | NA | NL |
| 182 | Galletitas dulces medialunitas | Ultra-Processed | Soybean | ARGENTINA | - | - | - | - | - | NA | NL |
| 183 | Mini *magdalena* cupcakes | Ultra-Processed | Soybean | FRANCIA | - | - | - | - | - | NA | NL |
| 184 | Sweet cookies with chocolate chips | Ultra-Processed | Soybean | ALEMANIA | - | - | - | - | - | NA | NL |
| 185 | Cookies with almonds and chocolate chips | Ultra-Processed | Soybean | FRANCIA | - | - | - | - | - | NA | NL |
| 186 | Sweet cookies decorated with chips | Ultra-Processed | Soybean | ARGENTINA | - | - | - | - | - | NA | NL |
| 187 | Sweet cookies with chocolate chips | Ultra-Processed | Soybean | URUGUAY | - | - | - | - | - | NA | NL |
| 188 | Sweet cookies with vanilla and chocolate flavor | Ultra-Processed | Soybean | URUGUAY | + | + | + | + | - | > 1% | NL |
| 189 | Lemon flavor waffle | Ultra-Processed | Soybean | BRASIL | - | - | - | - | - | NA | NL |
| 190 | American-style pancakes with *dulce de leche* | Ultra-Processed | Soybean | URUGUAY | - | - | - | - | - | NA | NL |
| 191 | Sweet cookies with vanilla and chocolate flavor | Ultra-Processed | Soybean | URUGUAY | + | + | + | - | - | > 1% | NL |
| 192 | Oatmeal sweet cookies with vanilla flavor | Ultra-Processed | Soybean | ARGENTINA | + | + | + | + | - | > 1% | NL |
| 193 | Assorted cookies | Ultra-Processed | Soybean | URUGUAY | - | - | - | - | - | NA | NL |
| 194 | Liquid soy food with fruit juice | Ultra-Processed | Soybean | ARGENTINA | - | - | - | - | - | NA | NL |
| 195 | Soy dietary food strawberry flavor | Ultra-Processed | Soybean | BRASIL | - | - | - | - | - | NA | NL |
| 196 | Liquid soy food sabor with vanilla flavor | Ultra-Processed | Soybean | ITALIA | + | + | - | + | - | >1% | NL |
| 197 | Whole-grain toast | Ultra-Processed | Soybean | BRASIL | - | - | - | - | - | NA | NL |
| 198 | Multigrain toasts | Ultra-Processed | Soybean | BRASIL | + | + | - | + | - | > 1% | NL |
| 199 | *Crockers* cookies with bran | Ultra-Processed | Soybean | ARGENTINA | + | + | + | - | - | > 1% | NL |
| 200 | Biscuit stick artificial flavor milk | Ultra-Processed | Soybean | BRASIL | + | + | + | - | - | <1% | NL |
| 201 | Salty rice cakes | Ultra-Processed | Soybean | ARGENTINA | - | - | - | - | - | NA | NL |
| 202 | Onion and parsley flavor cereal snacks | Ultra-Processed | Soybean | BRASIL | - | - | - | - | - | NA | NL |
| 203 | Instant soup with meat flavor | Ultra-Processed | Soybean | EEUU | + | + | + | + | - | > 1% | NL |
| 204 | Salty biscuit with bran | Ultra-Processed | Soybean | PARAGUAY | - | - | - | - | - | NA | NL |
| 205 | *Cracker* cookies | Ultra-Processed | Soybean | ARGENTINA | - | - | - | - | - | NA | NL |
| 206 | Rice crackers flavored with herbs | Ultra-Processed | Soybean | URUGUAY | + | + | + | - | - | > 1% | NL |

## Acknowledgments

The authors would like to acknowledge C.S.I.C. (Comisión Sectorial de Investigación Científica-UdelaR), Espacio Interdisciplinario-UdelaR, PEDECIBA, Biotechnology Master Program-UdelaR, and Intendencia de Montevideo for financial support of this work.

